# Optimisation of Xenium automated in situ sequencing for PAXgene-fixed tissue samples

**DOI:** 10.1101/2025.02.11.637091

**Authors:** Kenny Roberts, Andrew R. Bassett

## Abstract

Spatial transcriptomics has transformed the study of gene expression in tissue samples, yet its application to non-standard sample preservation formats beyond fresh frozen and FFPE remains underexplored. PAXgene fixation offers advantages for genomic studies by preserving DNA and RNA without crosslinking but poses challenges for high-quality spatial transcriptomics. Here, we present an optimised workflow for applying Xenium, an automated in situ sequencing platform, to PAXgene-fixed paraffin-embedded (PFPE) tissues. Using our newly developed Xenium Tissue Optimisation (XTO) protocol, we systematically tested permeabilisation conditions across multiple mouse and human tissues. We show that pepsin digestion effectively enhances RNA accessibility in PFPE samples, with tuneable digestion times for tissue-specific optimisation, though a compromise may need to be reached between transcript detection rates and maintaining tissue integrity and morphological staining. Our results indicate that, under optimal conditions, PFPE samples can yield comparable or superior spatial RNA transcript detection to formalin-fixed paraffin-embedded (FFPE) samples. This study provides a framework for adapting in situ sequencing to PFPE tissues, broadening the applicability of spatial transcriptomics to archival and prospective sample collections.

**Visual abstract:** 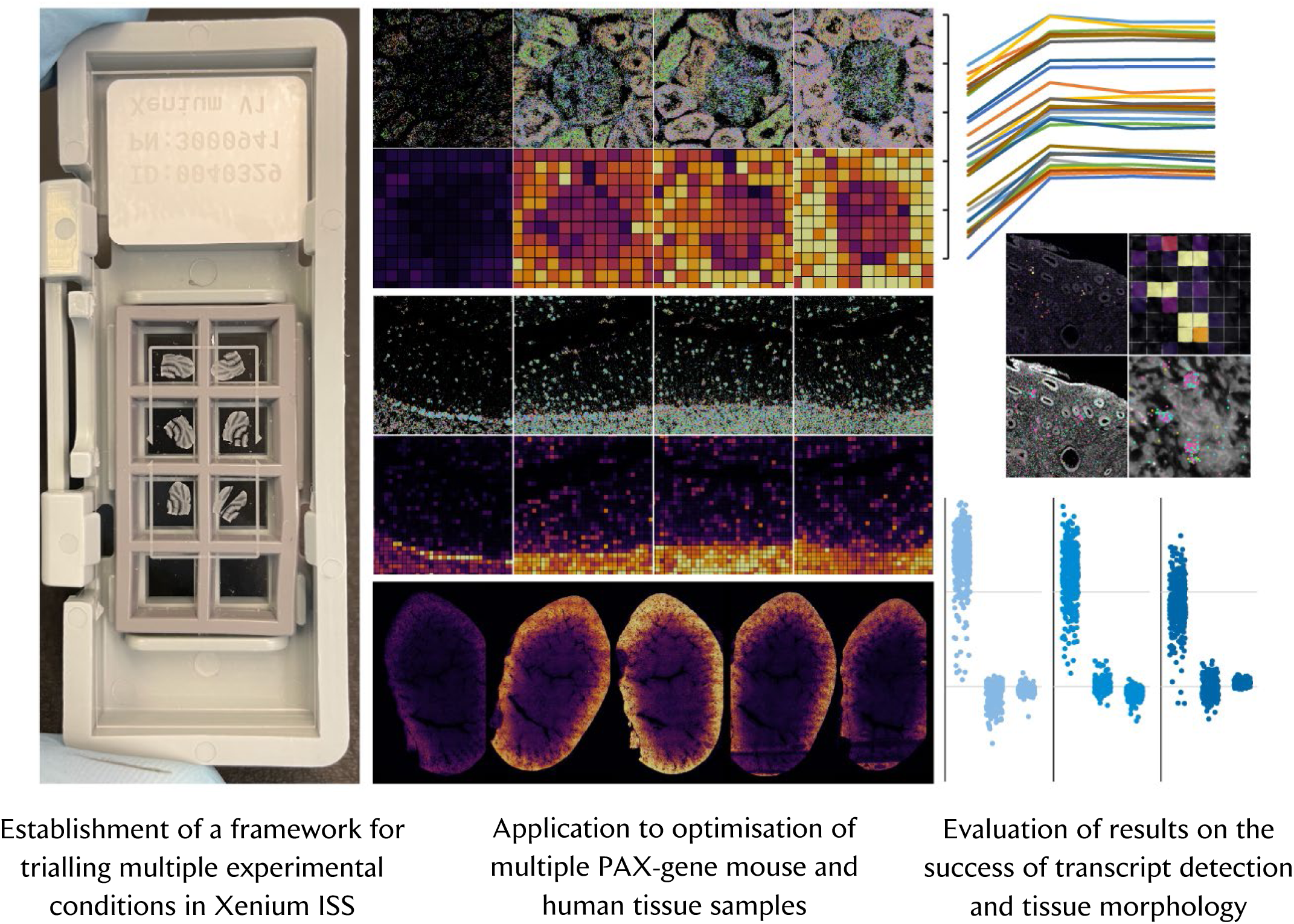

## Introduction

As the range of spatial transcriptomics technologies has exploded over the last five years, so has interest from scientific communities in applying these methodologies to diverse sample types. However, the use of archival biological material may be at odds with the preparation formats preferred by, or for which protocols are optimised by, technology providers.

The two most common preparations for mammalian (human, mouse) tissue samples are: so-called “fresh frozen”, where tissues are rapidly frozen using dry ice or liquid nitrogen; and FFPE (formalin-fixed paraffin-embedded), whereby tissues are chemically fixed using solutions of formaldehyde or paraformaldehyde prior to dehydration and embedding in paraffin wax. These and other preparations are summarised in Table 1. Processing routes are summarised in Figure 1.

**Figure 1:**
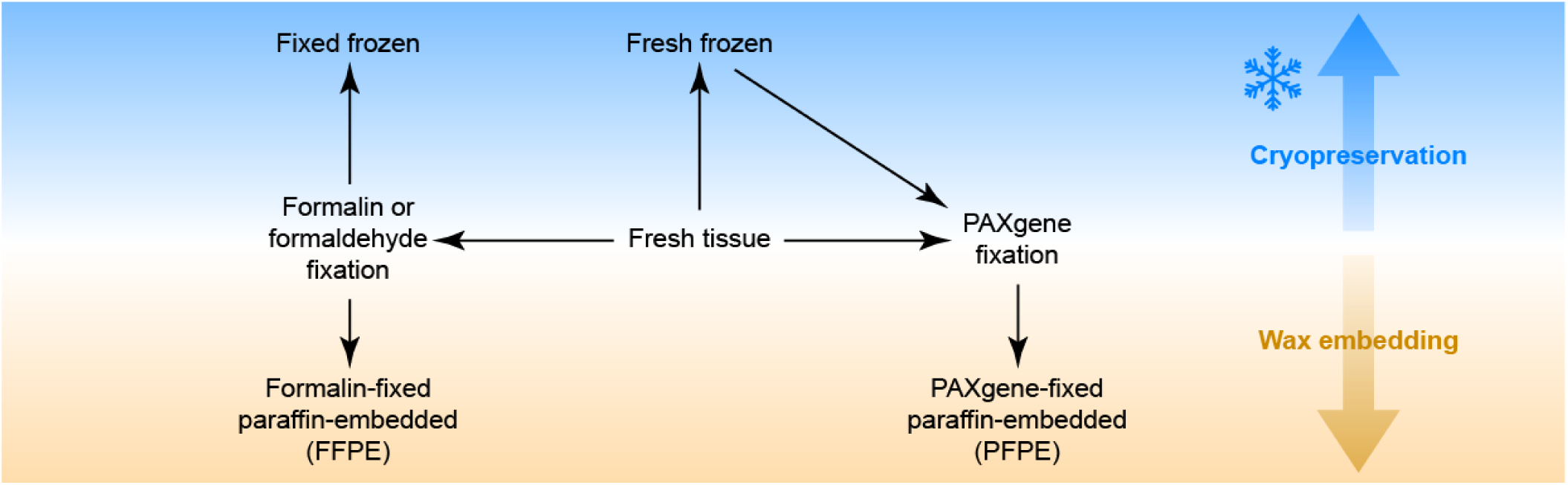
Sample processing routes for tissue samples for long-term storage.

**Table 1:** Different preparations of tissue samples for long-term storage.

| Preparation | Details | Usage details |
| --- | --- | --- |
| Fresh frozen<br>aka<br>'Snap frozen' | Tissue **frozen at approximately -80°C** using dry ice or liquid nitrogen. May be **embedded in 'optimal cutting temperature' (OCT) medium** beforehand. Samples stored at -80°C. | Morphological quality is dependent upon tissue and freezing conditions; poor freezing leads to irreversible artefacts. Good freezing and storage practices can maintain samples with high molecular integrity for years, including nucleic acids. High quality nucleic acids may be extracted and isolated, in addition to being targeted *in situ* by hybridisation-based approaches. Short fixation with formaldehyde solutions typically applied post-sectioning in order to preserve molecular integrity during staining protocols. |
| Fixed frozen | Tissue **fixed in formaldehyde or paraformaldehyde** solution, then **frozen** as per fresh frozen. Optionally, samples may be **cryopreserved** by impregnation with sucrose or glycerol solutions before freezing. | Commonly used for mouse necropsy tissue, as fixative may first be perfused transcardially. Has been inherited by some human brain labs. Short fixation may be used to toughen fragile human tissues prior to freezing. Depending upon duration of fixation, may be treated more similarly to fresh frozen or to FFPE (albeit excluding paraffin-related processing). |
| Formalin-fixed paraffin-embedded (FFPE) | Tissue **fixed using formalin** (formaldehyde with methanol as a stabiliser), typically for 16-24 hours or longer, then **dehydrated** using an ethanol series and cleared using xylene. Impregnated with molten paraffin wax and then **embedded in a wax block** that may be stored at 4-25°C. | A mainstay of pathology labs and histology services, long fixation yields excellent preservation of tissue morphology for downstream sectioning and H&E or histochemical staining. Dehydration and storage in wax preserves molecular integrity for long-term storage (decades if optimal). Storage at 4°C is recommended where the sample will be used for (spatial) transcriptomics. Nucleic acids and proteins are crosslinked, precluding optimal nucleic acid extraction, but a good substrate for probe hybridisation-based approaches following appropriate 'de-crosslinking' or epitope retrieval. |
| PAXgene-fixed paraffin-embedded (PFPE) | Tissue **fixed using PAXgene**, a proprietary alcoholic fixative, **then dehydrated and embedded as per FFPE**. | Offers the benefits of long-term storage of paraffin-embedded tissue but without the cross-linking nature of formaldehyde-based fixatives. PAXgene denatures and 'fixes' cells while leaving nucleic acids non-crosslinked and so amenable to extraction and isolation. |

Due to the prevalence and familiarity of fresh frozen and FFPE samples, these two preparations are almost exclusively favoured for the optimisation and validation of new single-cell and spatial genomics technologies. While this is logical, and the application of many technologies from the past decade to fresh frozen and FFPE tissues has yielded countless discoveries, the exclusion of other less widely used preparations limits the applicability of such technologies to other sample cohorts. The long-term storability that is a key intention of tissue preservation naturally leads to scenarios where samples were preserved in a particular manner before a new technology becomes available. One such preparation, which will be the subject of this manuscript, is PAXgene-fixed tissue.

PAXgene is a proprietary system (PreAnalytiX, QIAGEN) for chemical fixation that is marketed as an alternative to formaldehyde-based solutions such as formalin. Tissues are first fixed for 2-72 hours in PAXgene Tissue FIX, which according to the safety data sheet contains methanol (between 50 and 70% w/w) and acetic acid (between 10 and 20% w/w). Subsequently, tissues are directly exchanged into PAXgene Tissue STABILIZER for prolonged storage for days (room temperature), weeks (refrigerated), and years (frozen). The PreAnalytiX website advises that “the PAXgene Tissue STABILIZER contains 70% ethanol” and so this is analogous to storage after fixation with formalin. Following these steps, samples may be dehydrated and embedded in paraffin wax (PAXgene-fixed paraffin-embedded, PFPE) as per FFPE processing. Similarly to formalin fixation, the relative simplicity of this two-step process allows for samples to be preserved in challenging or time-pressured conditions (contrasting the more involved process of optimally cryopreserving tissue in OCT, for example). However, while formaldehyde cross-links and fragments nucleic acids, PAXgene does not – it is a ‘coagulative’ fixative – and thus it may be favoured for applications where high quality nucleic acids need to be extracted, such as for low-input DNA sequencing in combination with laser capture microdissection (LCM)^1^ or high-resolution single-molecule mutation mapping^2^.

To further complicate matters, PAXgene may be used as a post-freezing fixative, whereby snap frozen tissues are thawed and concurrently fixed in PAXgene Tissue FIX and then processed to paraffin. The PAXgene-fixed samples used in the experiments described hereafter were originally collected for genome sequencing following LCM and were originally frozen at the site of collection, whereafter they were thawed in PAXgene Tissue FIX and then processed through to PFPE.

It is not trivial to exhaustively optimise conditions for the staining of many different tissues, especially for a sample type such as PFPE where there is little existing literature. PFPE tissues have previously been shown to be amenable to DNA in situ hybridisation^3–5^ and also used to prepare samples for multiplexed smFISH using Resolve Biosciences Molecular Cartography^6–10^ but unfortunately the protocols are not publically available for the latter, including any pre-treatment or permeabilisation details. Nevertheless, in this study, a number of conditions were trialled for the preparation of PFPE slides for use with Xenium, an automated version of direct RNA-targeting hybridisation in situ sequencing (HybRISS)^11^.

## Results

Twenty pre-treatment regimes were screened to identify the best approach for PFPE tissue permeabilisation Initially a broad range of pre-treatment regimes were trialled, combining different fixation and epitope retrieval elements.

Three fixation regimes were trialled:

1. No additional fixation. Theoretically, PFPE tissues are already ‘fixed’ albeit not in a cross-linking fashion.
2. 30 minutes of post-fixation (after baking and de-paraffinisation) with 3.7% formaldehyde as per the recommended Xenium protocol for fresh frozen tissue. It was not clear whether this would be beneficial, perhaps from a perspective of RNase deactivation or additionally ‘fixing’ RNA in place once the samples were re-hydrated.
3. 24 hours of post-fixation with 3.7% formaldehyde in order to mimic an FFPE-like state. Prolonged fixation (8-12 hours) of fresh frozen tissue followed by FFPE-appropriate epitope retrieval has previously worked well in our hands for RNAscope on human decidua-placenta tissue^12^. This approach (24 hours) was also used previously for DNA ISH on PFPE human breast tissue^4^ and is the suggested protocol from the manufacturers of PAXgene for ISH assays (PreAnalytiX Supplementary Protocol PX21).

Following fixation (or not), four main types of epitope-retrieval regimes were trialled:

1. Permeabilisation with 1% SDS, as per the recommended Xenium protocol for fresh frozen tissue (the methanol incubation from this protocol was not included since PAXgene has methanol as a majority component).
2. Permeabilisation via digestion with pepsin. This was inspired by previous experience with CARTANA ISS^13,14^ as well as Spatial Transcriptomics, later 10x Genomics Visium.
3. Combined decrosslinking and protease digestion using the Xenium Decrosslinking protocol as per the recommended protocol for FFPE (and fixed frozen) tissue.
4. Decrosslinking in boiling citrate buffer. This was inspired by previous experience with CARTANA ISS for fixed frozen and FFPE tissues.

Not all fixation and epitope-retrieval combinations were trialled, as some are illogical e.g. decrosslinking in citrate buffer or Xenium decrosslinking reagent is not necessary where there has been no significant fixation with formaldehyde. In all, twenty conditions were devised and trialled in the first experiment (Fig. 2A) including a ‘no pre-treatment’ control. At this stage, in order to be able to screen multiple tissue types and so many pre-treatment regimes, Xenium was not used as an output. Instead, following pre-treatment, slides were stained with RNAscope smFISH, allowing large batches of slides to be stained in parallel in an automated fashion on a Leica BOND RX. Slides were stained with a 3-plex mouse ‘positive control’ panel of broadly expressed housekeeping genes including the highly expressed *Ubc* (ubiquitin C). While the RNAscope and Xenium protocols rely on different types of amplification, the probe lengths are similar (∼40 nt of target-binding region, divided in both cases into two co-dependent ∼20 nt sequences) and the hybridisation temperatures are only 10°C different (40°C for RNAscope, 50°C for Xenium).

**Figure 2:**
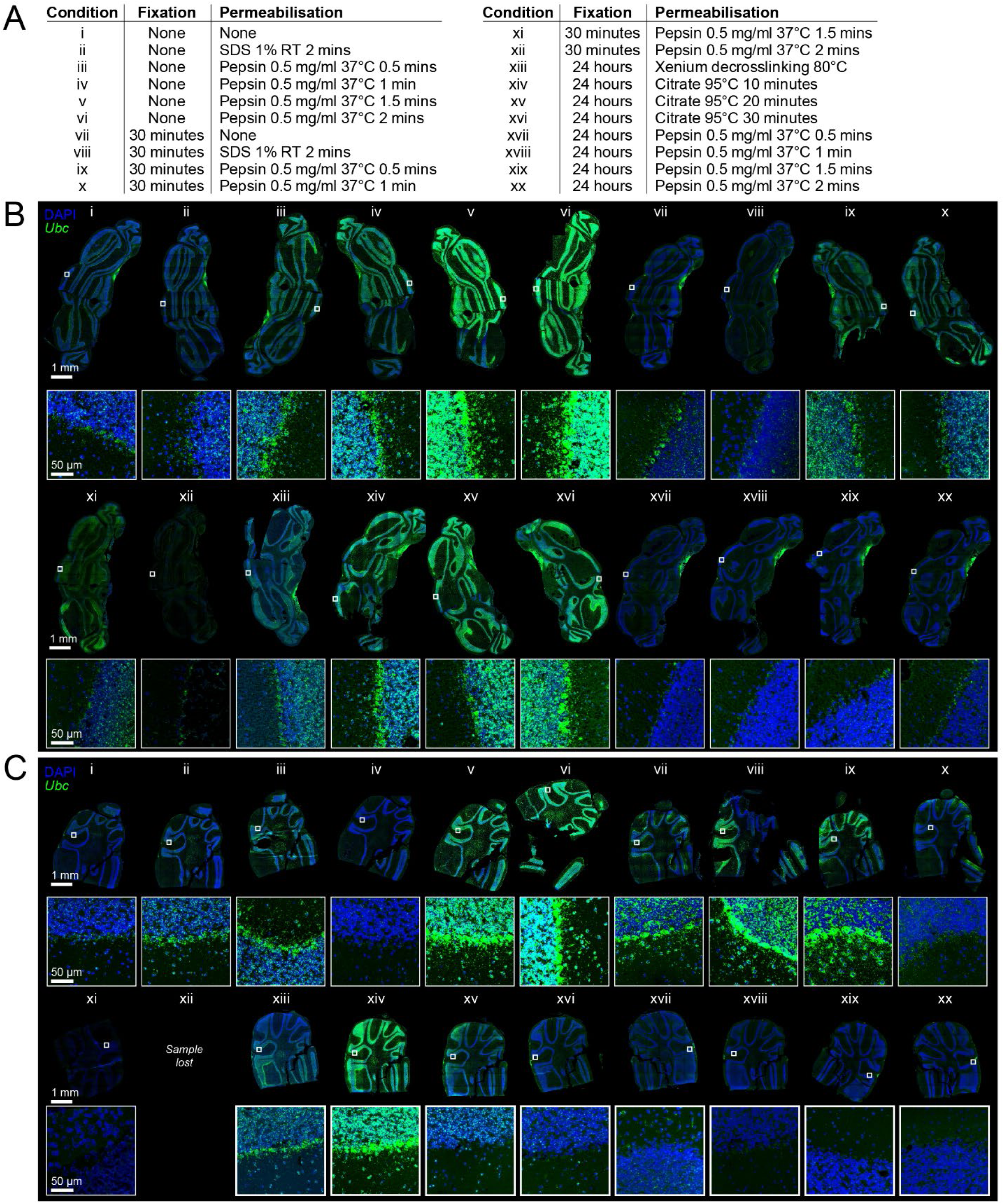
Narrowing down potential pre-treatment approaches using RNAscope smFISH as a proxy readout. (A) A total of 20 pre- treatment regimes were trialled using an RNAscope smFISH readout. (B-C) RNAscope staining of replicate sections of two mouse cerebellum tissue samples (from different animals) following aforementioned pre-treatments. *Ubc* RNA staining shown in green, DAPI-stained nuclei in blue. All full-tissue section images shown to the same scale (scale bar 1 mm); all inset images shown to the same scale (scale bar 50 µm).

Of the different ‘categories’ of pre-treatment trialled, protease digestion with pepsin without prior post-fixation yielded the highest *Ubc* signal intensity across both samples (Fig. 2B-C, conditions v and vi), and was ultimately chosen for further experiments. Long post-fixation followed by boiling in citrate buffer also performed well (Fig. 2B-C, conditions xiv-xvi) and illustrates that there is not always a single route to an experimental end, but pepsin digestion gave higher *Ubc* signal intensities, is less likely to result in tissue detachment, and is easier to integrate into the existing Xenium workflow in a streamlined way retaining use of the gasket-on-a-thermocycler architecture. Examining the other two genes in the RNAscope panel – *Polr2a* (RNA polymerase II subunit A) and *Ppib* (peptidylprolyl isomerase B) – actually revealed that citrate buffer yielded better signals for one of the samples (Supp. Fig. 1A). Two other tissues, kidney and liver, were also stained following a range of pepsin digestion times, and illustrated that good staining can be obtained from multiple tissues using pepsin, but that individual tissues may require different digestion times for optimal signal (Supp. Fig. 1B-C).

A Xenium Tissue Optimisation (XTO) protocol was developed to optimise Xenium performance on PFPE tissues In order to make most economical use of Xenium reagents and runs, a method was implemented to trial multiple conditions on the same Xenium slide, by combining it with the gasket architecture from the 10x Genomics Visium Tissue Optimisation (TO) protocol to create a Xenium Tissue Optimisation (XTO) experiment. Tissue sections were placed on the Xenium slides so as to align within both the Xenium capture area and the capture areas created by the rubber seal of the Visium TO gasket (Fig. 3A-B). A maximum of six of the eight wells created by the Visium TO gasket can be aligned with the Xenium sample area (Fig. 3A). Following baking and de-paraffinisation, slides were briefly dried at 37°C then sealed into Visium TO gaskets and subjected to different digestion times (Fig. 3C-G). Depending upon the tissue section size, the rubber gaskets were in some cases cut using a scalpel in order to combine sample areas to create larger ‘wells’ (Fig. 3C).

**Figure 3:**
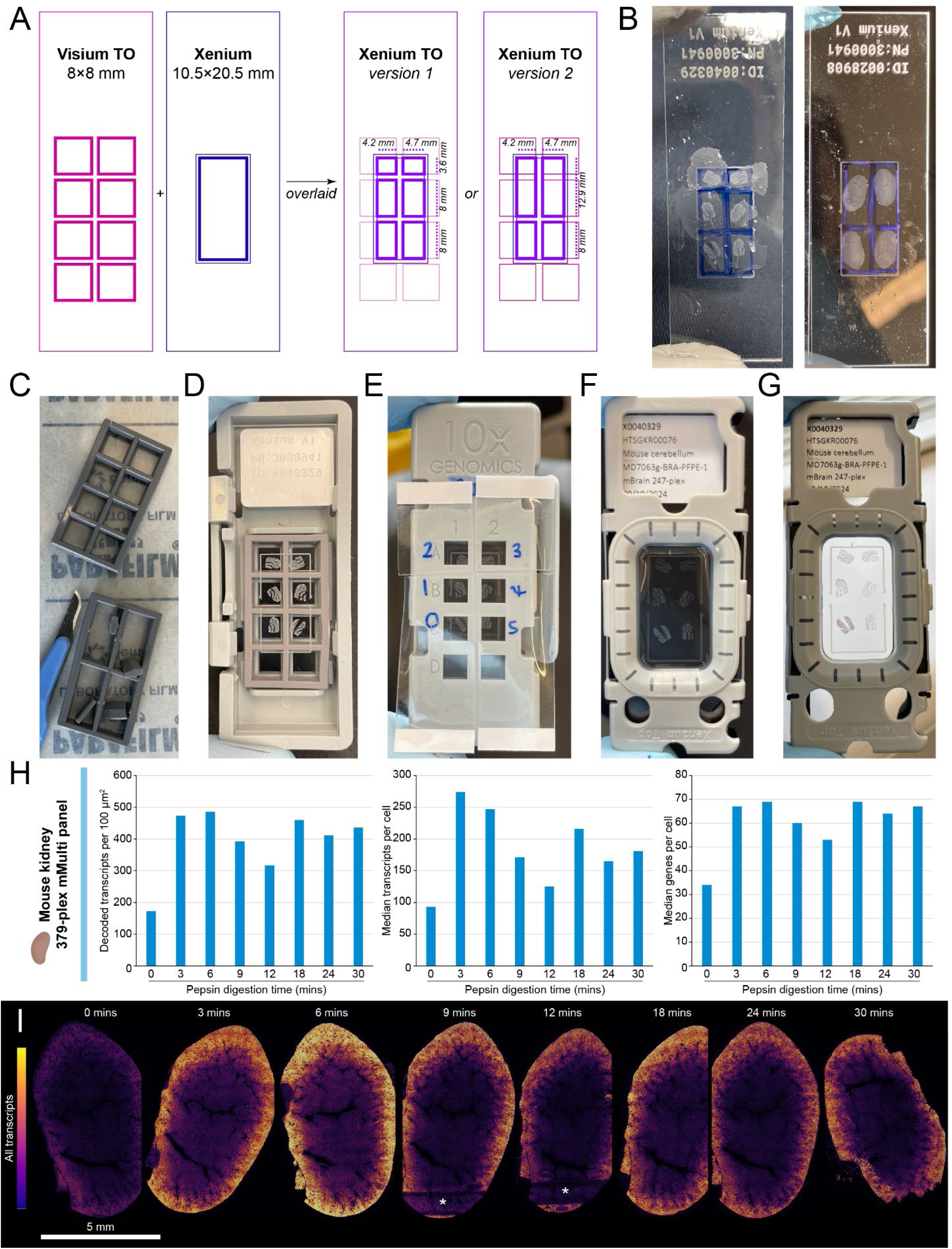
Implementation of a Xenium Tissue Optimisation (XTO) protocol. (A) Overlaying the layouts of the Visium TO and Xenium slides results in six areas of common sample area (version 1) which can be partly combined into four larger areas (version 2). (B) Paraffin sections are placed upon Xenium slides so as to align with these common areas (drawn for reference on the back of the slide with a marker pen); mouse cerebellum (left) and mouse kidney (right) shown. Following optional cutting of the rubber Visium TO gasket (C), slides are assembled into the Visium TO cassette (D, bottom view) such that one section is in each well, allowing reagents to be added to individual sections via the openings on the top of the cassette (E, top view, labelled for pepsin digestion time course). Progressive digestion of sections is visible by differential opacity immediately following digestion (F), and later during autofluorescence quenching (G). (H) Xenium transcript and gene count metrics and (I) spatially resolved transcript density in PFPE mouse kidney following time course of pepsin digestion using the XTO protocol. *N.B.* The areas of low density and disrupted morphology (asterisks) result from sections falling outside the usable area and being in contact with the rubber gasket.

For the first experiment, eight replicate sections of a PFPE mouse kidney sample were processed across two slides with a time course of pepsin digestion and then assayed with the 379-plex 10x Genomics pre-designed Xenium Mouse Tissue Atlassing Panel (mMulti). The range of pepsin digestion timings was similar to those recommended in the Visium TO kit, ranging from 0 to 30 minutes (though the pepsin activity in the latter is not exactly known, the activity used here is inspired by the original Spatial Transcriptomics protocol^15^). It was anticipated that the typical optimal digestion time for PFPE for Xenium would be substantially less than that for fresh frozen tissues for Visium since the latter relies upon sufficient digestion that RNA is able to diffuse through the tissue and be captured by the coating on the Visium slide while successful Xenium ISS requires only that the RNA molecules are accessible to probes but fundamentally still localised within the cell.

The optimal digestion time in this preliminary experiment was between 0 and 6 minutes, with the number of transcripts detected per cell and per unit area (100 µm^2^) peaking at 3 minutes and 6 minutes respectively (2.9-fold and 2.7-fold increases over the no digestion, respectively) (Fig. 3H). These numbers are consistent with the overall transcript density across the sections, which peaked visually at 6 minutes (Fig. 3I). Following a decrease in transcripts and genes from 6 to 12 minutes, the metrics unexpectedly increased at 18-30 minutes (Fig. 3H), though not higher than the initial peak at 3 to 6 minutes. However, these extended digestion times led to progressive loss of nuclear morphology (Supp. Fig. 2A).

Five genes in the 379-plex panel were detected more highly in the undigested section: *Chl1* (cell adhesion molecule L1-like), *Ube2c* (ubiquitin conjugating enzyme E2 C), *Ncf4* (neutrophil cytosolic factor 4), *Upk3a* (uroplakin 3A), and *Scg2* (secretogranin II). All of these show broad and weak expression across the undigested sample with no overt cell type specificity (Supp. Fig. 2B), though this does not explain their sudden drop-off following pepsin digestion. Detection of the other 374 genes increased following pepsin digestion, 367 by at least 2-fold and 142 of which by at least 5-fold (Supp. Fig. 2C).

Across all transcripts detected, one gene was substantially more highly detected at 3 minutes than 6 minutes: *Cox8b* (cytochrome c oxidase subunit 8B) is a thermoregulatory Complex IV subunit associated with adipose cells in the perirenal fat^16^ (Supp. Fig. 2D). This hinted that the same permeabilisation conditions may not suit both adipose and the body of the kidney. Accordingly, examining the density of the mostly highly detected genes in a region of perirenal adipose including *Cox8b*, the adipogenesis regulator *Car3* (carbonic anhydrase 3)^17^, the gluconeogenesis control enzyme *Pck1* (phosphoenolpyruvate carboxykinase 1), and fatty acid transporter *Cd36*^18^ illustrates that this region of tissue was optimally permeabilised at 3 minutes (Supp. Fig. 2D), with total transcript density increasing by 6.3 fold from 0 to 3 minutes of pepsin and then decreasing by the same fold from 3 to 6 minutes (Supp. Fig. 2E-F). This is divergent from the overall section transcript count and density which is mostly stable at 3 versus 6 minutes (Supp. Fig. 2E-F).

This experiment was then repeated with a second mouse kidney sample (the other organ from the same mouse). Six replicate sections were digested for between 0 and 6 minutes in order to examine the transcript detection curves with greater granularity (Fig. 4A). In parallel, six replicate sections of a mouse cerebellum sample were assayed with the 247-plex 10x Genomics pre-designed Mouse Brain Gene Expression Panel (mBrain) following a digestion time course from 0 to 5 minutes (Fig. 4B). It was anticipated based upon the nature of the two organs and also the previous experiments with RNAscope as an output that the optimal digestion time for cerebellum would be shorter than that for kidney.

**Figure 4:**
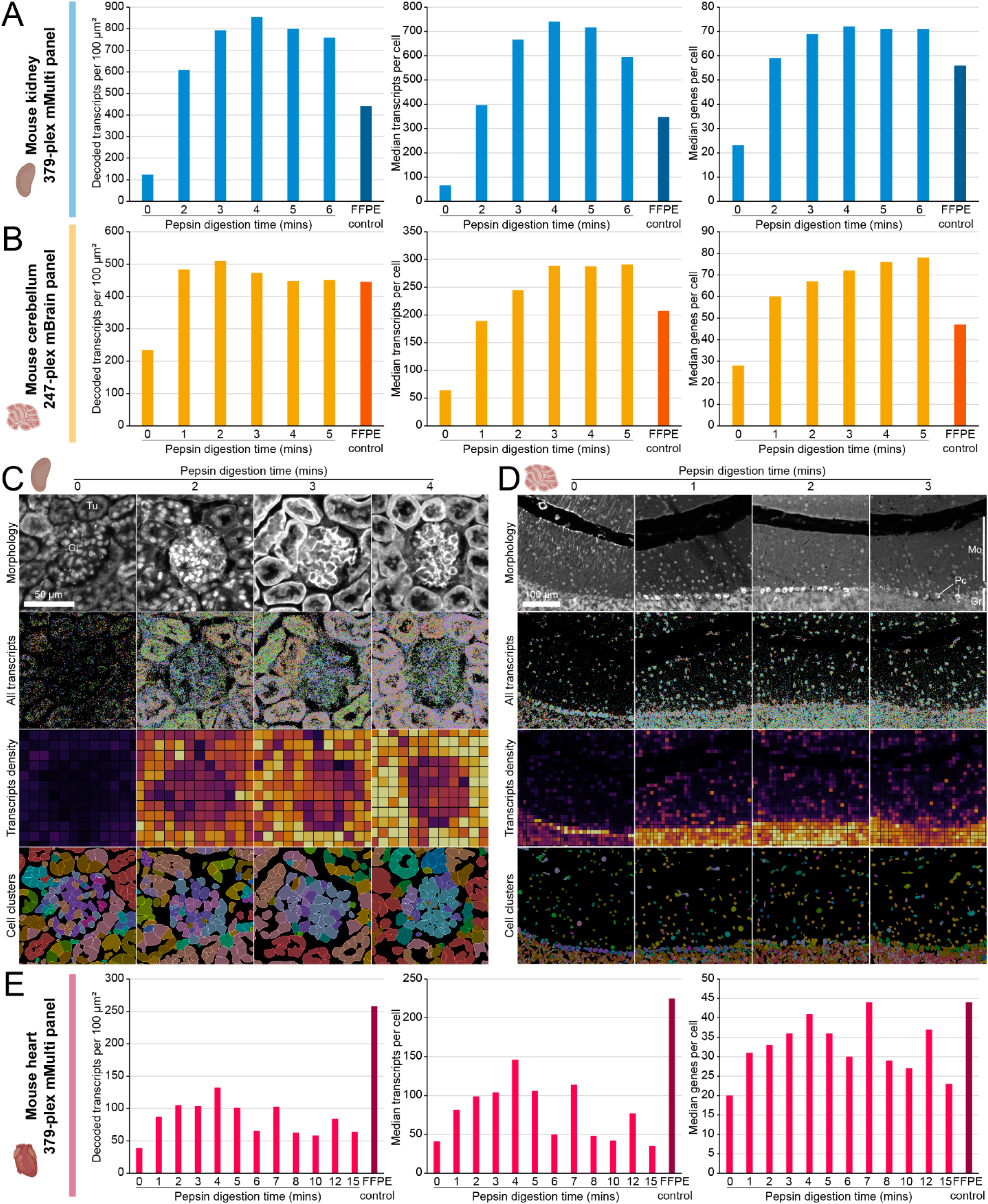
Application of the Xenium Tissue Optimisation (XTO) protocol to various PFPE mouse tissues. (A-B) Xenium transcript and gene metrics obtained from: (A) application of a 6 minute XTO time course to PFPE mouse kidney subsequently stained with the mMulti panel; and (B) application of a 5 minute XTO time course to PFPE mouse cerebellum subsequently stained with the mBrain panel. FFPE comparison sample metrics shown to the right of each plot, darker colours. (C-D) Segmentation stain image, transcript detection, and cell type clustering for (C) PFPE mouse kidney (Gl = glomerulus, Tu = tubule) and (D) PFPE mouse cerebellum (Gr = granular layer, Mo = molecular layer, PC = Purkinje cell) across XTO permeabilisation. (E) Xenium transcript and gene metrics for PFPE mouse heart tissue following 15 minute XTO permeabilisation and assay with the mMulti panel. FFPE sample metrics shown to the right of each plot, darker colours.

Both experiments gave smooth curves of Xenium transcript detection metrics across the range of permeabilisation. Consistent with the previous experiment that indicated an optimal permeabilisation time around 3-6 minutes, kidney yielded maximal transcripts both per cell and per unit area with 4 minutes of pepsin digestion (6.9-fold and 11.4-fold increases over no digestion, respectively) (Fig. 4A). A maximal number of transcripts per cell in cerebellum was reached at 3 minutes (3.8-fold over no digestion) and plateaued thereafter. The number of transcripts per 100 µm^2^ for cerebellum actually peaked at 2 minutes (Fig. 4B), indicating that apparent cell size is altered by increasing pepsin treatment. Indeed the average number of cells per unit area decreased for both kidney and cerebellum with increasing digestion (Supp. Fig. 3A). Nuclear transcript detection rates were similar to whole transcript detection rates (Supp. Fig. 3B). For both kidney and cerebellum, detection of all transcripts was increased with permeabilisation relative to the untreated section. Permeabilisation unveiled greater transcript detection across different cell types and anatomical features, including glomeruli and tubules of the kidney (Fig. 4C) and the granular and molecular layers of the cerebellum (Fig. 4D).

Broadly, non-specific signals as assessed using the Xenium ‘adjusted negative control probe rate’, which is calculated using proprietary probes that should not bind to any transcripts in the tissue, increased as samples were permeabilised with pepsin, but particularly beyond the point of optimal transcript detection (Supp. Fig. 3C). All values were in line with typical negative control probe rates for mouse and human tissues in our experience.

To provide a comparison for transcript metrics, FFPE mouse kidney and liver tissues were prepared and assayed according to the standard 10x Genomics protocols for FFPE tissue. The results are shown alongside those for the aforementioned PFPE samples (Fig. 4A-B). For both kidney and cerebellum, transcript metrics for optimal PFPE treatment were higher than the comparable FFPE sample, suggesting that RNA integrity and targetability in the PFPE sample is indeed higher.

One additional PFPE mouse tissue was assayed, in order to attempt to define an upper bound for the length of prospective permeabilisation experiments. Heart is one of the toughest tissues; human heart requires some of the longest digestion times for Visium, up to 45 minutes^19^, compared to typically 18-20 minutes for kidney, for example. To allow for a potentially much higher degree of permeabilisation, twelve replicate sections of PFPE mouse heart were processed across two XTO slides with a pepsin time course covering up to 15 minutes of digestion. Unexpectedly, maximal transcript detection occurred after 4 minutes of pepsin treatment, the same as for mouse kidney (Fig. 4E). Note that metrics for 7 minutes and 12 minutes time points were considered outliers due to inconsistent section thickness, which can be observed on the XTO slide following permeabilisation and following autofluorescence quenching (Supp. Fig. 3D). While the transcript per unit area and per cell metrics for this PFPE heart sample fell short of the FFPE comparison sample, the maximum number of genes per cell was similar, indicating that even sub-optimal transcript detection has a limited impact upon the diversity of gene expression visible (Fig. 4E).

Intriguingly, of the five genes whose detection density decreased in the heart from 0 to 4 minutes of permeabilisation, four overlap with the five whose detection fell during permeabilisation of the kidney (*Chl1*, *Ube2c*, *Ncf4*, and *Scg2*) and the remaining one of the latter *Upk3a* is the gene with the second lowest increase in density (only 1.09-fold) (Supp. Fig. 3E). This suggests a specific relationship between permeabilisation and the accessibility of these transcripts or behaviour of the probes targeting them. If the physical transcripts for these genes or the cells that express them are especially friable, they may be lost with even mild permeabilisation but, as in the kidney (Supp. Fig. 2B), these genes do not appear to be well expressed or overtly cell type-specific in the heart (Supp. Fig. 3F). Alternatively, the probes targeting these transcripts may be particularly prone to off-target binding in undigested PFPE tissue, and digestion may reduce this noise. It is notable that the negative control probe rates in both the kidney and heart drop following permeabilisation (Supp. Fig. 3C).

An interesting observation was the impact of permeabilisation on the performance of the Xenium In Situ Cell Segmentation Kit. This is a cocktail of antibodies and probes designed to facilitate better cell segmentation by staining cell membrane proteins as well as RNA and proteins within the cell. The antibody stains are somewhat cell type-dependent, owing to differences in expression of the target proteins. However, the ‘interior RNA’ stain, which targets ubiquitous 18S ribosomal RNA, has performed well in our hands across many different tissues and cell types. Since it targets RNA, it is logical that the performance of this component is dependent upon RNA accessibility in the same way as mRNA targets for ISS. Accordingly the intensity of the interior RNA stain was broadly correlated with time of permeabilisation and the density of transcripts detected in the three mouse tissues (Supp. Fig. 3G-I), though was generally lower in the heart. Maximum intensities in PFPE samples were similar to or lower than those observed in the FFPE comparison samples (Supp. Fig. 3G-H). The efficacies of the ‘membrane’ (ATP1A1, E-cadherin, and CD45) and ‘interior protein’ (ɑ-smooth muscle actin and vimentin) antibody components were also affected by pepsin digestion, but with differential effects per tissue. For example, in the non-permeabilised cerebellum, the interior protein stain (presumably the vimentin antibody) marks surface blood vessels that descend into the molecular layer, but this staining was lost with even mild pepsin digestion (Supp. Fig. 3G), while it was preserved excellently in the FFPE comparison sample. In the kidney, on the other hand, vimentin staining of podocytes in glomeruli and also of tubules appeared only *after* pepsin digestion (Supp. Fig. 3H). Staining intensity with the boundary antibody mix increased after pepsin treatment in both cerebellum and kidney (Supp. Fig. 3G-H), though its utility in actually defining cell edges in the former is limited. In the heart, the interior protein stain, presumably the ɑ-smooth muscle actin antibody, stained smooth muscle around blood vessels in the untreated section, but this staining disappeared with permeabilisation (Supp. Fig. 3I). Boundary antibody staining increased with permeabilisation, and can actually be seen to mark sarcomeres in cardiomyocytes captured in the longitudinal orientation (Supp. Fig. 3I, second row insets).

### The XTO protocol allows rapid optimisation of Xenium for PFPE human tissue samples

As a proof of principle that the approach detailed in this work may be applied to human patient-derived samples, and also to investigate the applicability of a single condition to multiple samples of the same tissue from different donors, the protocol was applied to optimise application of Xenium to PFPE human uterus from donors of reproductive age, with a focus on the endometrium. Three uterine tissue samples containing endometrium and myometrium were each sectioned onto a 6 capture area XTO slide and permeabilised across a 5 minute time course, then processed with the 377-plex 10x Genomics pre-designed Xenium Human Multi-Tissue and Cancer Panel (hMulti). The resulting data are also discussed in greater detail as a ‘worked example’ of choosing the optimal permeabilisation condition for a given tissue.

The three samples each yielded curves of transcript detection metrics broadly similar to those seen in the aforementioned mouse tissue experiments. Surveying whole-section metrics, transcript and gene detection appeared to peak following between 1 and 3 minutes of pepsin digestion (Fig. 5A) indicating some differences between samples, though all demonstrated excellent improvements over the untreated section (14.1- to 22.8-fold increases in transcript density; 11.8- to 26.7-fold increases in median transcripts per cell). As in mouse tissues, the nuclear transcript detection rates strongly mirrored the overall transcript detection (Supp. Fig. 4A, left), and cell density per area trended downwards with permeabilisation (Supp. Fig. 4A, centre). Interestingly, the negative control probe detection rates dropped dramatically following any permeabilisation (4.5- to 13.1-fold decreases) (Supp. Fig. 4A, right), though the values remain relatively high. Adjusted negative control probe rates of 0.02 and 0.04 are approximately at the 80th and 90th percentiles, respectively, based on our collective Xenium experiment history (approximately 450 tissue sections over 150 slides of many different FFPE and fresh frozen tissues).

**Figure 5:**
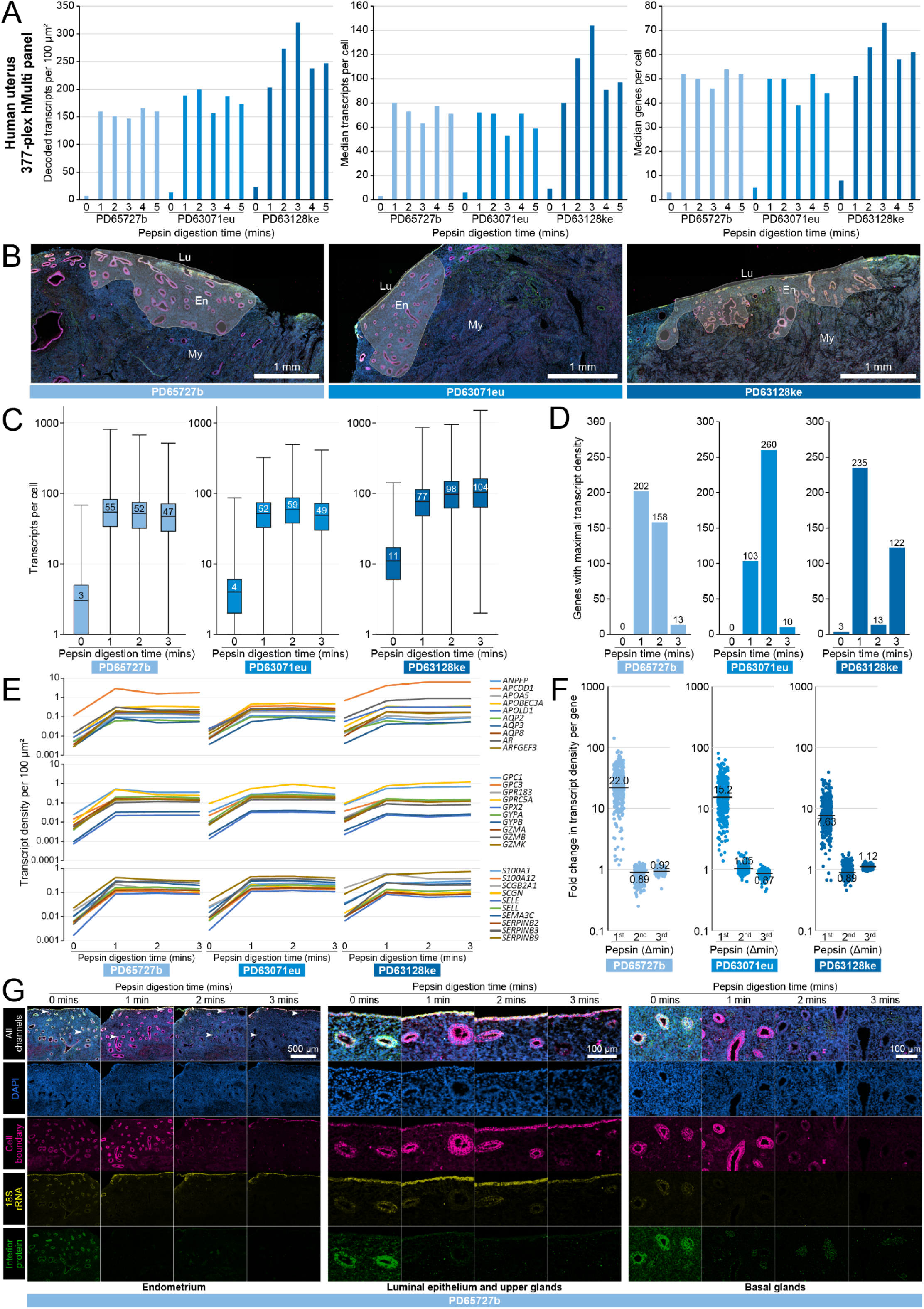
Application of the Xenium Tissue Optimisation (XTO) protocol to human uterus tissue. (A) Xenium transcript and gene detection metrics for three PFPE uterus samples (three biological replicates from different donors PD…, colour coded shades of blue throughout this figure) assayed with the human multi-tissue and cancer (hMulti) panel following a 5 minute XTO time course. (B) Selection of endometrium-specific regions for further analyses (Lu = lumen, En = endometrium, My = myometrium). (C) Distribution of transcript counts per cell in endometrium across the XTO time course. Central lines depict medians, boxes represent interquartile ranges, whiskers depict ranges. (D) Numbers of genes detected maximally at each time point of the XTO duration. (E) Example curves of transcript detection for 30 alphabetically selected genes (10 each above, middle, below) for three samples across XTO permeabilisation. (F) Incremental fold changes in transcript detection for all genes across three samples. (G) Segmentation stain images of one representative donor, showing large tissue area (left, scale bar 500 µm) and magnified regions of upper and lower glandular areas (middle and right, scale bars 100 µm), indicated at top-left with white arrowheads.

It was anticipated that the endometrium and myometrium tissues may behave differently, with the muscle-rich myometrium likely being more resilient to permeabilisation than the more fragile endometrium. Regions of endometrial tissue were selected in each sample (Fig. 5B), based upon the observation of luminal and glandular epithelia and the absence of: a) strong expression of *DES* (desmin) and *MYH11* (myosin heavy chain 11), muscle-specific genes that designate the myometrium; and b) any artefacts resulting from the gasket placement in the XTO protocol (Supp. Fig. 4B). Total transcript counts per cell for the selected endometrial region were plotted per time point (Fig. 5C), as well as the number of genes for which transcript detection peaked in each condition (Fig. 5D). Differences remained in the time point at which transcript detection peaked per sample, indicating that this did not result from the conflation of different proportions of endometrium and myometrium tissue in the overall section metrics. In PD64727b and PD63071eu, the peak of median transcripts per cell coincided with the time point at which the maximum number of genes exhibited a maximal density of transcript detection, at 1 minute and 2 minutes, respectively (Fig. 5C-D, left and middle). PD63128ke demonstrated a peak of genes with maximum density at 1 minute, but the median number of transcripts per cell peaked at 3 minutes, coinciding with a secondary peak in the number of genes with maximum detection density (Fig. 5C-D, right). Examining neither these, nor plots of individual gene transcript detection profiles (some are shown in Fig. 5E), gave a firm conclusion as to the optimal condition, the latter also illustrating that gene expression levels intrinsically vary widely across the panel (2-3 orders of magnitude per sample) and together indicating that total transcript count is prone to bias.

A metric was sought to identify the ‘best’ condition by: a) focusing upon the specific tuneable parameter of permeabilisation, i.e. incremental time of incubation; and b) using an overall parameter that is more robust to the range of expression levels of each gene. The effect that each incremental step in permeabilisation had on detection of each gene was calculated as the geometric mean of individual gene transcript density fold changes for each n^th^ minute of permeabilisation. Across the endometrium regions of the three samples, the first minute of pepsin digestion had by far the greatest impact on transcript detection (22.0-fold, 15.3-fold, and 7.6-fold, for the three samples) (Fig. 5F), while the second and third minute of digestion time changed transcript density only by between 0.88- and 1.11-fold, with the detection of many genes being negatively impacted by the second and/or third minute (Fig. 5F) in spite of the overall transcript density increase in PD63071eu and PD63128ke (Fig. 5C).

Only a few genes in one sample – *COL17A1* (collagen type XVII alpha 1 chain), *MZB1* (marginal zone B and B1 cell specific protein) and *TMEM52B* (transmembrane protein 52B) in PD63128ke – were negatively impacted by the first minute of permeabilisation (Fig. 5F, right), and these genes exhibit some of the lowest density fold changes in the two samples where they are positively affected by digestion. Plotting the collective expression density of the six genes common to the list of ten with the lowest fold change in the first minute of permeabilisation in each sample shows that they are enriched in the glandular epithelium (Supp. Fig. 4C), suggesting that either it is among the less durable compartments of this tissue; and/or it is more prone to non-specific binding to the probes of these genes.

Interestingly, on a finer level the detected expression pattern of *MZB1* changes more dramatically than the detected expression level: with no permeabilisation, *MZB1* is detected broadly and weakly throughout the tissue, with some enrichment in glands as above; however, upon permeabilisation, expression strongly peaks within sparse cells (Supp. Fig. 4D, top), where it is co-expressed with other established B-cell markers including *CD79*, *CD27*, and *TENT5C* (terminal nucleotidyltransferase 5C) (Supp. Fig. 4D, bottom), indicating that the detected signals are more reflective of biological expression in situ following permeabilisation.

Finally, the morphological quality of the tissue sections was evaluated across the permeabilisation time course (Fig. 5G). In the untreated section, all of the segmentation stain components (boundary, interior RNA, and interior protein) marked epithelial cell populations. The 18S rRNA stain showed some selection for the luminal epithelium, and this was retained as the tissue was permeabilised, though glandular staining was significantly weakened. With no permeabilisation, the boundary and interior protein cocktails showed some differential staining of gland populations (Supp. Fig. 4E), with the upper (functional) glands having a higher ratio of the interior protein stain, which also stained the stroma in a somewhat non-specific fashion. This was lost following permeabilisation, which strongly reduced glandular and stromal interior protein staining, restricting it to blood vessels (Fig. 5G, bottom rows), while the intensity of the boundary stain was increased in glandular epithelium (Fig. 5G, middle rows) during the first minute of permeabilisation. After 2 minutes, tissue morphology was negatively impacted, particularly in the case of glandular epithelium which showed loss of nuclear clarity and cell membrane staining (Fig. 5G, Supp. Fig. 4F).

Considering both the transcript metrics and the segmentation stain images, we would conclude an optimal permeabilisation time of 1 minute for this endometrium tissue cohort. While some samples may deliver higher transcript densities with longer permeabilisation times, this comes at the expense of tissue morphology and the introduction of more bias in terms of inter-sample variability of individual gene expression profiles.

## Discussion

We have shown here for the first time successful staining of RNA using in situ sequencing in PFPE samples, demonstrating that it is possible to generate high-quality state-of-the-art spatial transcriptomics data from sample cohorts processed using this alternative fixative. Although the comparison is limited by the fact that the samples were prepared independently from different animals and demonstrate some differences in anatomical profiling, our observation that PFPE samples can yield higher transcript counts than FFPE samples is: a) consistent with previous reports that RNA quality is higher in PFPE samples^20–22;^ and b) promising for the application of Xenium and other spatial genomics protocols to existing PFPE sample cohorts. It is worth noting however that prolonged storage of PFPE blocks has been reported to result in RNA degradation that brings the quality to FFPE levels or below^23^. We observed higher transcript counts in FFPE mouse heart than in PFPE but, although we cannot rule out a tissue-specific effect, it is possible that the PFPE heart sample was prepared sub-optimally upstream. Further analyses are needed to establish whether PFPE is a recommendable format for prospective sample collection; it could be an ideal format for preserving tissue for parallel genomic and transcriptomic analyses.

We have explored here several approaches to the identification of the ‘best’ condition for a given sample/cohort. Compared to the Visium Tissue Optimisation experiment that inspired the XTO protocol, where evaluation typically consists of simply identifying the brightest tissue section, the breadth of data generated by Xenium allows for a deeply comprehensive evaluation incorporating both transcript and morphological parameters on the scale of single-cells, tissue features, and whole sections. However, this also makes the conclusion more complicated. Ideally, we would take advantage of the range of metrics provided directly by the Xenium instrument such as the transcript density or count per cell. Since the assignment of transcripts to cells is a key element of biological interpretation of spatial transcriptomics data, the transcripts *per cell* metric seems more relevant than transcripts per 100 µm^2^ when selecting which condition is optimal. However, this is highly dependent upon the accuracy and biological meaningfulness of what is deemed by the Xenium Analyzer software as a cell. Even beyond the inherently challenging nature of accurate cell segmentation, we have shown the impact that pepsin treatment may have on nuclear morphology as well as cell segmentation stains, both of which impact segmentation accuracy. We have not ventured here into post-hoc cell segmentation, which further complicates analysis but may prove a worthwhile investment over a longer time scale. The change in transcript density per incremental time of permeabilisation gave a more unifying conclusion across the three human endometrium samples assayed here than the transcripts per area or cell metrics per sample. It may be necessary to compromise between total transcript detection and tissue integrity, where nuclei start to be damaged at longer digestion times. It is also important to focus the XTO evaluation on regions or sub-tissues of interest to ensure that the optimal conditions are selected for the prospective experiment.

Ultimately, we recommend that any XTO experiments are evaluated holistically, incorporating not only the various transcript and gene metrics but also negative control rates, segmentation accuracy, and expression of known marker genes. Although the 10x Xenium multi-tissue and cancer panel is intended for use on a wide range of tissues, it lacks key markers for the endometrium and it would be interesting to assess whether the use of a more tailored gene panel would a) affect the optimal condition; or b) make it easier to identify by providing better signposts of expression in particular tissues or cells of interest.

While we only ran Xenium v1 here (i.e. up to 480-plex), the optimisation should be applicable to the Xenium Prime (5000-plex) assay too, since the pre-treatment regimes for the two assays are identical for FFPE and for fresh frozen tissues, with the exception of a shorter bake for FFPE tissues for the Xenium Prime assay (presumably to minimise any RNA degradation). However, this needs to be tested empirically as the chemistry of the assays does differ.

Optimised conditions for mouse tissue samples are not necessarily representative of human tissue samples. Samples were not available to optimise conditions for the same organ in both species in order to test this empirically, though it is encouraging that the human endometrium samples required a similar time to multiple mouse tissues. Broadly we expect most tissues to be made amenable to Xenium by some degree of pepsin permeabilisation, though some may require a different strategy not explored here. Only mouse cerebellum tissue was screened with the full range of 20 conditions and used to select pepsin as the permeabilisation agent for further testing, though the results from other tissues shown here are very promising. It is possible that very tough tissues such as human heart may require a longer pepsin time course in spite of the mouse heart results presented here, and that unusual tissues such as bone or eye may require a broader assessment of conditions such as different enzymes.

With the samples available for this study, we were not able to address whether prior freezing of samples before fixation with PAXgene affects the overall quality or the optimal permeabilisation conditions for Xenium downstream but it seems likely that the overall strategy would be compatible regardless. Furthermore, while non-standard sample preparations other than PFPE were not addressed here, elements of the approach would be applicable. A pepsin time course may be effective for the optimisation of ethanol-fixed paraffin embedded (EFPE) tissue, for example, while the use of the Visium TO gasket to segregate the Xenium slide into multiple independent regions could be useful for other treatments. There is wide interest in using spatial technologies such as Xenium to assay non-mammalian cells such as bacteria, and these experiments would also benefit from the ability to trial many permeabilisation conditions in a cost- and time-effective manner. The use of the Visium TO cassettes does require an additional purchase and add an additional cost to the Xenium experiments, approximately 25% extra per XTO slide, though the cassettes and gaskets are reusable (so long as they produce a good seal) and this would reduce the extra cost.

The concept of using RNAscope as a proxy to identify the best permeabilisation conditions for a given tissue is appealing for several reasons, including cost and throughput. However, the differential performance between the three genes presented here across conditions suggests that effective use of RNAscope to determine optimal Xenium conditions would rely upon identification of specific genes that are most indicative of Xenium performance across the range of permeabilisation, and this may be tissue-, disease state- and cohort-specific. The reason for lack of correlation between *Ubc* and the other two RNAscope targets under some conditions is not clear. Likewise the reason that a small minority of genes in the mouse kidney and human endometrium Xenium experiments dropped out following even the shortest pepsin permeabilisation, but any potential reasons – differential permeabilisation of subcellular compartments, differential dissociation of multi-molecular complexes, etc. – are only speculative.

This study establishes a framework for adapting spatial transcriptomics protocols to diverse tissue preservation methods, specifically demonstrating the feasibility of optimising Xenium for PFPE samples. Expanding our approach to other spatial technologies could further enhance the utility of archival and non-traditional tissue samples in generating insightful multi-omic datasets.

## Methods

### Human tissue sample ethics

Human uterus tissue was collected from donors of reproductive age at Addenbrooke’s Hospital, Cambridge University Hospitals (15/EE/0152, East of England–Cambridge South Research Ethics Committee).

### PFPE and FFPE tissue block preparation

For PFPE samples, mouse tissues and human tissues were snap frozen on dry ice and stored temporarily at -80°C. Samples were then thawed in PAXgene Tissue FIX (PreAnalytiX) and fixed for 24 hours, then transferred to PAXgene Tissue STABILIZER for 16-24 hours.

For FFPE samples, mouse tissues were fixed in 10% neutral buffered formalin for approximately 24 hours and then serially dehydrated in 50%, 70%, and 100% ethanol.

Fixed, dehydrated samples were cleared with xylene and impregnated with paraffin wax using a Tissue-Tek VIP 6 Tissue Processor, then embedded into cassettes.

### Microtome sectioning

PFPE and FFPE sections were cut at 5 µm using a Leica RM2265 Automated Microtome.

Placement of multiple paraffin sections in such close proximity for the XTO protocol is not trivial and great care must be taken when the slide is in the water bath in order to prevent adjacent sections detaching from the slide. It may be necessary to experiment with atypical angles or even immersing the ‘top’ end of the slide for the collection of the first sections.

### Pre-treatments

Fixation: a 3.7% formaldehyde solution was prepared by ten-fold dilution of 37% formaldehyde solution (Fisher 10532955) with 1× PBS, as per the 10x Genomics protocol for fresh frozen Xenium (CG000581). After incubation in 3.7% formaldehyde at room temperature for 30 minutes or 24 hours, slides were washed and paused in 1× PBS prior to other pre-treatments.

SDS permeabilisation: a 1% SDS solution was prepared by ten-fold dilution of 10% SDS solution (Sigma 71736) with nuclease-free water (Thermo AM9922), as per the 10x Genomics protocol for fresh frozen Xenium (CG000581). No methanol incubation was performed, slides were instead rinsed and paused in 1× PBS to await RNAscope staining.

Epitope retrieval using the 10x Genomics Xenium decrosslinking mix: urea and a permeabilisation enzyme were diluted with a decrosslinking buffer, and applied to the slide in a Xenium cassette as per the 10x Genomics protocol for FFPE Xenium (CG000580). Slides were washed with PBS-Tw (1× PBS with 0.05% Tween-20) prior to Xenium or RNAscope staining.

Pepsin digestion: a 1 mg/ml solution of pepsin was produced by dissolving pepsin powder (≥250 units/mg, Sigma P7000) in 0.1 M HCl (Fisher 10325710), yielding a solution of activity ∼250 units/ml.

For initial RNAscope screening, pepsin solution was pre-warmed in 50 ml Falcon tubes in a 37°C water bath, and slides were incubated in these tubes.

For the XTO protocol, aliquots of 1.5 ml of pepsin solution in 1.7 ml Eppendorf tubes were pre-warmed in a 37°C heating block. Solution was then pipetted into XTO reaction chambers. Afterwards, slides were immediately quenched in several washes of 1× PBS at room temperature.

Note: for the initial RNAscope trials, a different pepsin product was used (≥2,500 units/mg, Sigma P7012). This is a more concentrated form of the enzyme and is provided as lyophilised flakes that are extremely static-prone and more difficult to dissolve thoroughly. The higher activity per unit mass makes the product more prone to autolysis and we have anecdotally found that activities of resulting solutions of P7012 are prone to be: a) less consistent; and b) not proportional in activity to solutions prepared with P7000.

Citrate buffer epitope retrieval: citrate buffer was produced by ten-fold dilution of Citrate Buffer, pH 6.0, 10×, Antigen Retriever (Sigma C9999) with nuclease-free water (Thermo AM9922). 250 ml of 1× citrate buffer in a Tissue-Tek Slide Staining Dish was heated to 99°C using a Braun steamer, and slides were immersed using a Tissue-Tek Slide Holder for 10-30 minutes with constant heating. Slides were immediately quenched in several washes of 1× PBS at room temperature.

### Xenium Tissue Optimisation (XTO)

A detailed protocol for implementing the Xenium Tissue Optimisation protocol from sectioning to data may be found here: dx.doi.org/10.17504/protocols.io.n92ldr819g5b/v1.

Briefly, PFPE sections were placed onto Xenium slides and then dried, baked at 42°C for 3 hours, and stored as per the 10x Genomics protocol for FFPE Xenium sectioning (CG000578). Slides were deparaffinised as per the 10x Genomics protocol for FFPE Xenium (CG000580); briefly, slides were baked at 60°C and then incubated in xylene and rehydrated through an ethanol series and finally water. The only deviation from the protocol was that the slides were baked at 60°C for 30 minutes instead of 2 hours (as per the Xenium Prime protocol for FFPE).

Xenium slides were assembled into Visium Tissue Optimisation cassettes and gaskets. A permeabilisation time course was conducted by adding pepsin solution to individual reaction chambers and then quenching with PBS. Slides were removed from the Visium cassettes and assembled into Xenium cassettes.

### Xenium in situ sequencing

Following pre-treatments or the XTO protocol, probe hybridisation and all following steps were performed as per the 10x Genomics Xenium protocol (CG000582), with temperature-regulated steps performed on a C1000 Touch Thermal Cycler (Bio-Rad).

Slides were loaded onto the Xenium Analyzer as per the 10x Genomics Xenium protocol (CG000584). Mouse tissue datasets were generated using Xenium instrument software version 3.1.0.0 or 3.1.3.1, both with analysis version xenium-3.1.0.4. Human uterus tissue datasets were generated using Xenium instrument software version 3.2.1.2, with analysis version xenium-3.2.0.7.

Xenium datasets were viewed and interrogated using Xenium Explorer 3 (10x Genomics), which was also used to generate images and transcript density maps.

### RNAscope smFISH

Following manual pre-treatments, slides were stained using a Leica BOND RX to automate staining with the RNAscope Multiplex Fluorescent Reagent Kit v2 Assay (Advanced Cell Diagnostics, Bio-Techne) using the RNAscope LS mouse positive control probe set 321818, according to the manufacturers’ instructions. Prior to loading, slides were dehydrated through a series of 50%, 70%, 100%, and 100% ethanol, for 5 minutes each. No additional epitope retrieval was performed on the instrument but a dummy protease step was included in order to quench endogenous peroxidase activity with hydrogen peroxide. Tyramide signal amplification with Opal 520, Opal 570, and Opal 650 (Akoya Biosciences) was used to develop three RNAscope probe channels (*Polr2a*, Opal 650; *Ppib*, Opal 570; *Ubc*, Opal 520).

### Confocal imaging

Slides stained with RNAscope were imaged using a Perkin Elmer Opera Phenix Plus High-Content Screening System. Imaging was performed in confocal mode with 2 μm z-step distance, using a 40× (NA 1.1, 0.149 μm/pixel) water-immersion objective. Channels: DAPI (excitation 375 nm, emission 435–480 nm), Opal 520 (excitation 488 nm, emission 500–550 nm), Opal 570 (excitation 561 nm, emission 570–630 nm), Opal 650 (excitation 640 nm, emission 650–760 nm). Confocal image stacks were stitched as two-dimensional maximum intensity projections using proprietary Acapella scripts provided by Perkin Elmer. Images were visualised using OMERO Plus (Glencoe Software).

All RNAscope images are visualised with no gamma adjustment, meaning that visual intensity is representative of quantitative RNA signal intensity.

## Author contributions

K.R. and A.R.B. conceived the study. K.R. designed, performed, analysed, and interpreted experiments and wrote the manuscript, with input from A.R.B.

## Acknowledgements and funding

PFPE mouse tissue samples were kindly provided by Alex Cagan and Adrian Baez Ortega, and were originally sourced from Linda Partridge at UCL. FFPE mouse tissue samples were originally sourced from the Wellcome Sanger Institute Research Support Facility and processed with help from Liz Tuck. PFPE human endometrium samples were kindly provided by Luke Wylie and Peter Campbell, and were originally collected by Kourosh Saeb-Parsy and processed by Yvette Hooks.

Thank you to Liz Tuck, Luke Wylie, Yvette Hooks, and Lucy Yates for helpful discussions, and Andrew Trinh for assistance with preliminary work. Thank you to the Wellcome Sanger Institute Sequencing Operations team for assistance with Xenium runs. 10x Genomics provided reagents for assaying six slides of Xenium, with kind thanks to Mike Schnall-Levin. This work was funded by an institutional grant 220540/Z/20/A awarded to the Wellcome Sanger Institute.

---

**Supplementary Figure 1:**
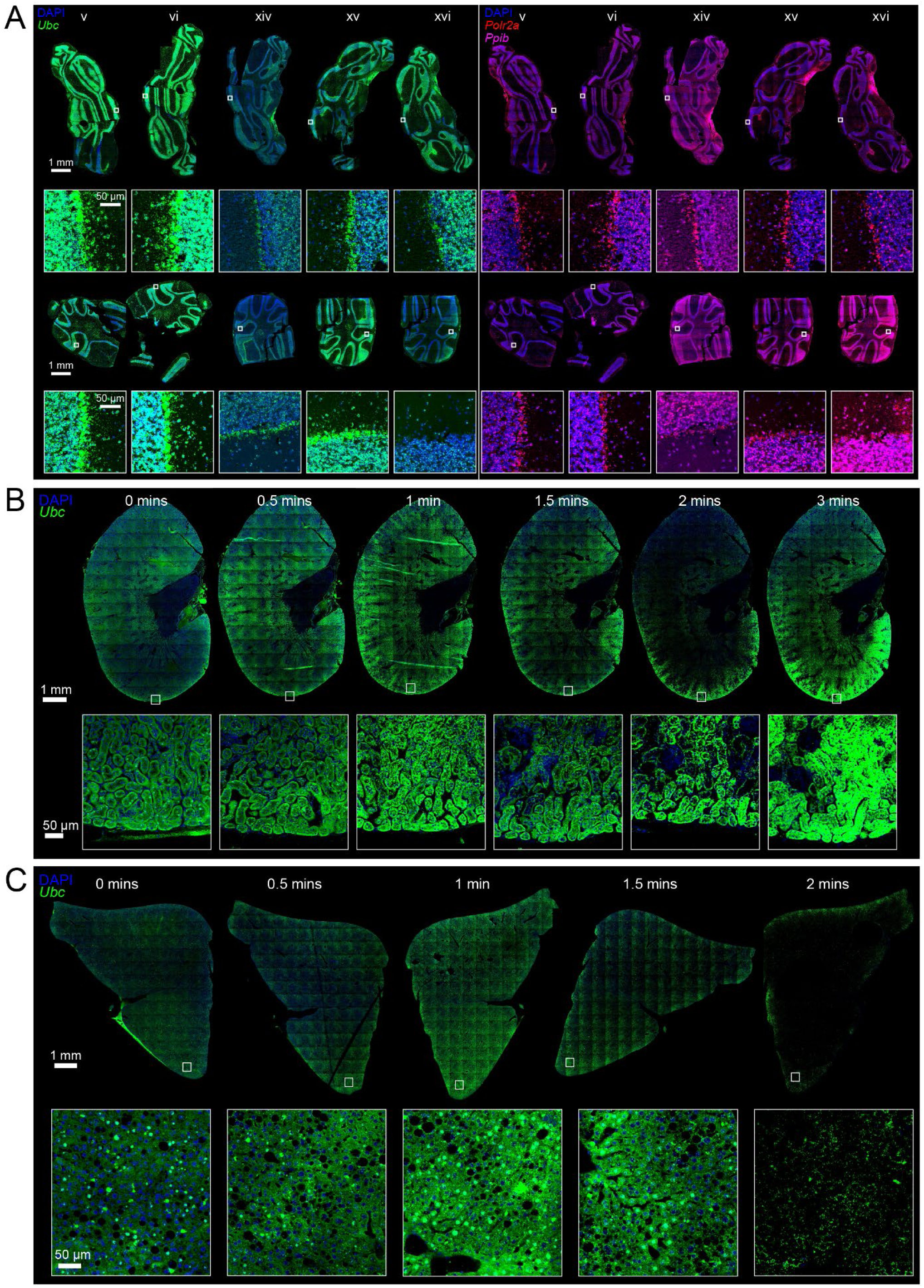
Different tissues yield optimal RNAscope smFISH signals after different durations of pepsin digestion. (A) Comparison of RNAscope smFISH staining for *Ubc* (left, green) versus *Polr2a* and *Ppib* (right, red and magenta) for the same five replicate sections of two cerebellum samples as are shown in Fig. 2B-C. All full-tissue images shown to the same scale (scale bar 1 mm); all inset images shown to the same scale (scale bar 50 µm). Samples with the brightest *Ubc* staining do not necessarily have the brightest *Polr2a* and *Ppib* staining. (B-C) RNAscope smFISH staining for *Ubc* following pepsin digestion time courses of PFPE mouse kidney (B) and PFPE mouse liver (C).

**Supplementary Figure 2:**
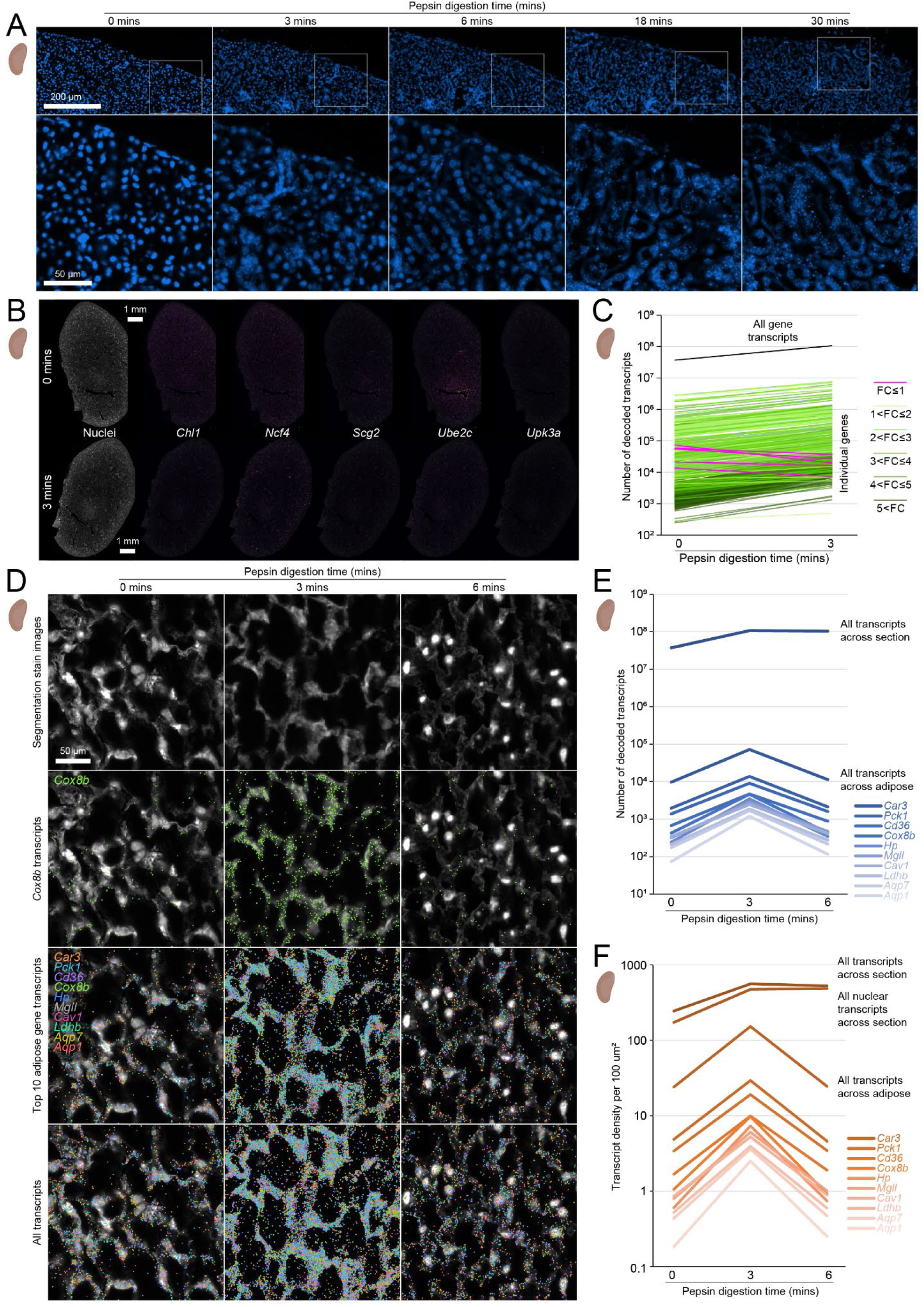
Impact of permeabilisation on PFPE mouse kidney nuclei and adipose cells. (A) DAPI-stained nuclei in PFPE mouse kidney across 30 minute XTO time course. Top row scale bar 200 µm, boxes indicate regions shown at higher magnification below, scale bar 50 µm. (B) Transcript density for five genes negatively impacted by permeabilisation, scale bar 1 mm. (C) Total number of decoded transcripts across entire tissue sections at two time points, shown per gene (colours) and for all genes (black). (D) Cell images (top) and transcript densities (*Cox8b*, top 10 expressed transcripts in adipose, and all transcripts, in respective rows) for adipose cells in perirenal fat across XTO time points. Scale bar 50 µm. (E-F) Total number of decoded transcripts (E), and average transcript density per 100 µm^2^ (F), for adipose-specific and all gene transcripts across both selected adipose tissue regions and entire sections.

**Supplementary Figure 3:**
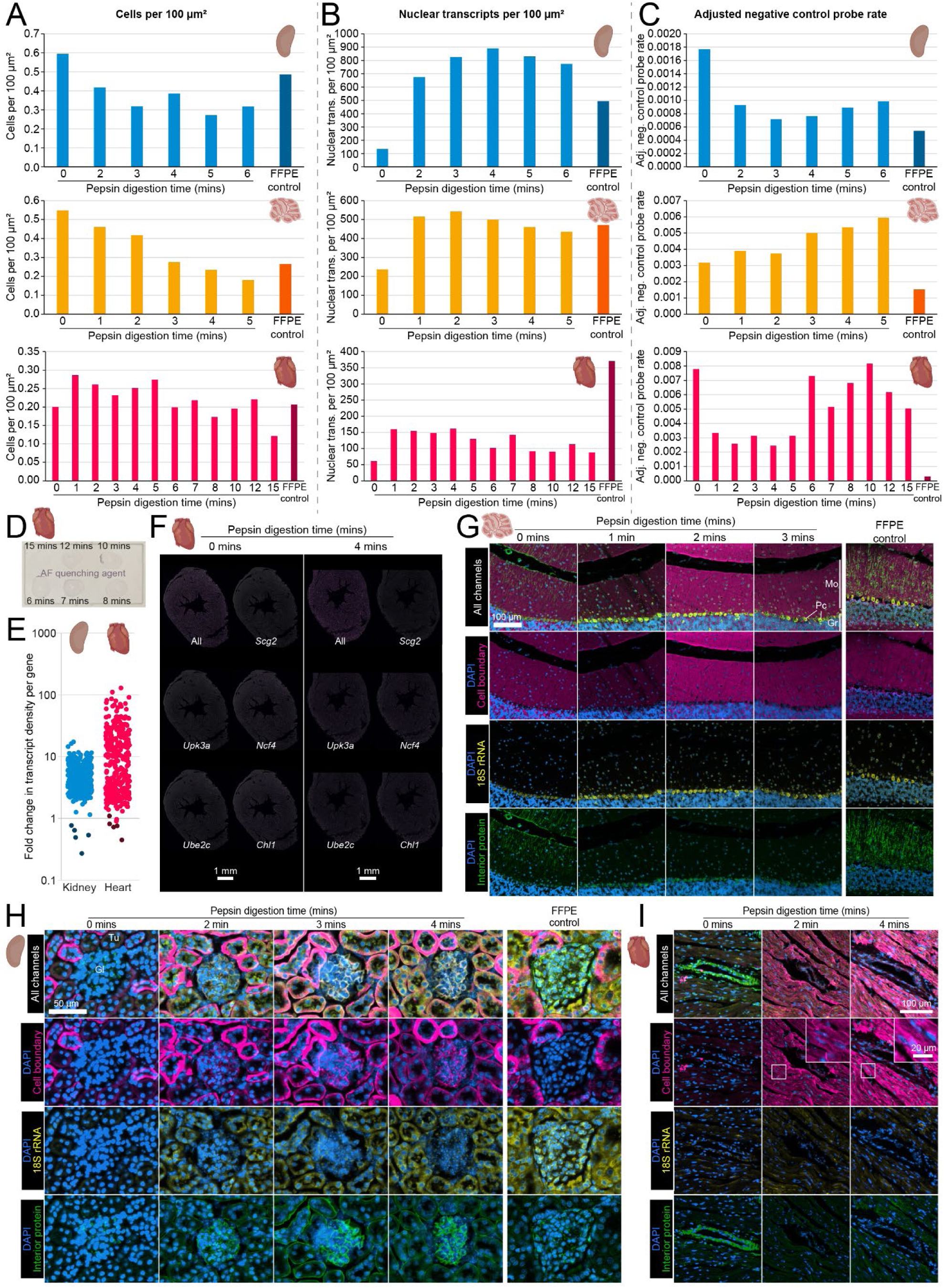
Further considerations for the evaluation of the XTO protocol in PFPE mouse tissues. (A-C) Additional Xenium metrics for the PFPE mouse tissue XTO pilot experiments including cell density (A), nuclear transcript detection (B), and negative control probe rates (C). (D) Photograph of one PFPE mouse heart XTO slide during Xenium processing, at the stage of drying post-addition of autofluorescence quenching agent, which stains the tissue sections purple. It can be seen that the 7 minute and 12 minute sections appear darker than expected, indicating that they are thicker than expected and explaining the anomalous transcript results shown in Fig. 4E. (E) Fold changes in transcript detection over the first increment of XTO permeabilisation in PFPE mouse kidney (Fig. 3H-I) and heart (Fig. 4E). The five genes with the lowest fold changes in kidney are coloured darker for both tissues. (F) Transcript density in heart for the same five genes, scale bar 1 mm. See also Supp. Fig. 2B. (G-I) Segmentation stain images for PFPE mouse cerebellum (G, scale bar 100 µm), kidney (H, scale bar 50 µm), and heart (I, scale bar 100 µm and inset scale bar 20 µm), across the XTO time course.

**Supplementary Figure 4:**
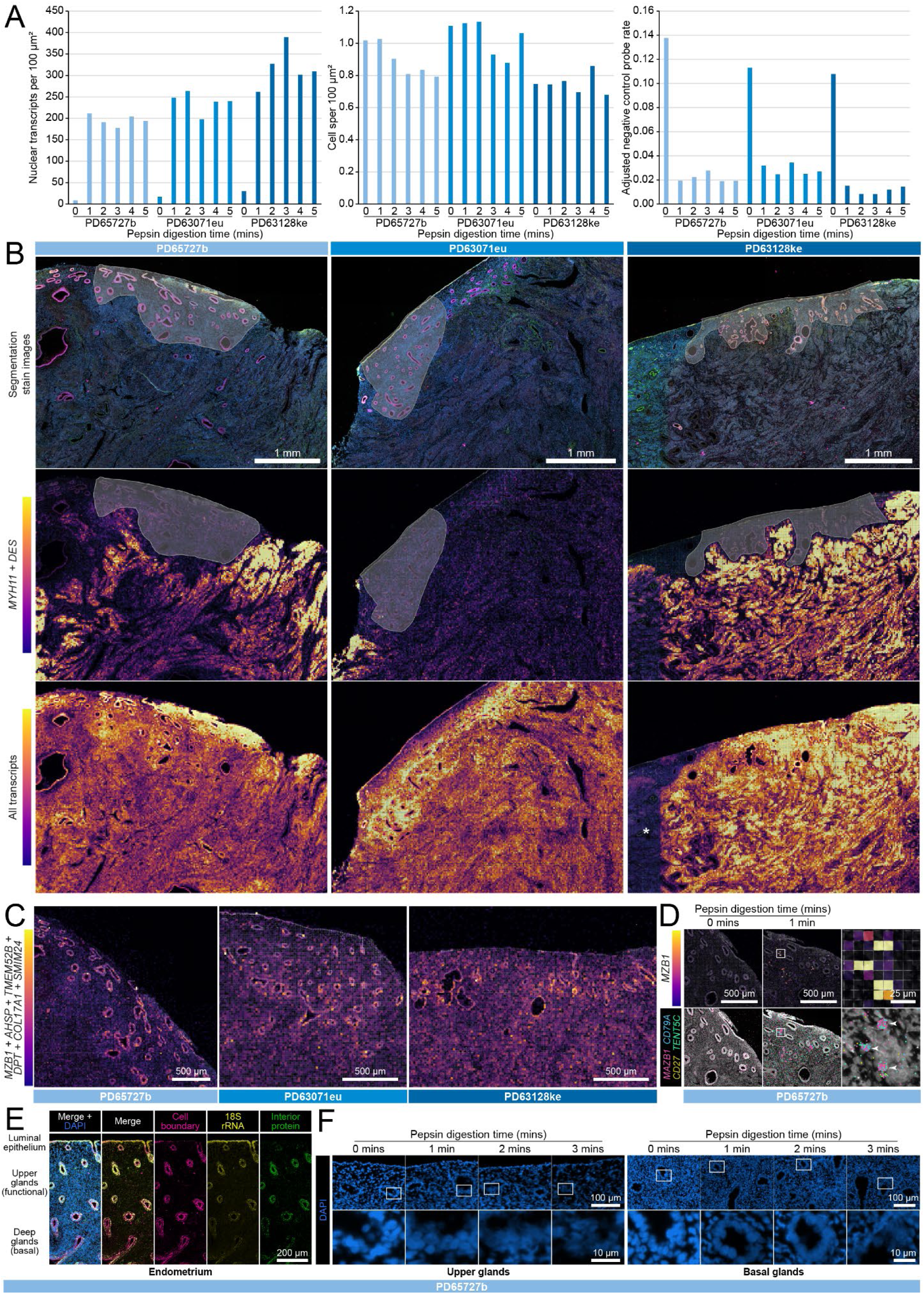
Further considerations for the evaluation of the XTO protocol in PFPE human uterus tissue. (A) Additional Xenium metrics of transcript and cell detection, and negative control probe rates for PFPE human uterus. (B) Selection of endometrium regions based upon glandular epithelium (above) and excluding regions of high *DES* and *MYH11* myometrium tissue (middle) or regions of low overall transcript density (below, see asterisk in bottom right panel for example). (C) Glandular epithelium enrichment of transcript detection of most friable genes across all three samples. (D) Transcript density for *MZB1* (top) and individual transcripts of *MZB1* and other B-cell markers (below) in non-permeabilised (left) and permeabilised (middle) endometrium. Area of high *MZB1* density is shown magnified to the right, where arrowheads highlight B-cells. (E) Differential staining of glandular epithelial populations with the segmentation staining kit, one representative donor shown. (F) Loss of nuclear morphology over XTO time course in epithelium from upper (left) and lower (right) glands. Scale bars 100 µm (above) and 10 µm (below), magnified regions indicated by boxes above.

